# Extinction probabilities of small structured populations: adequate short-term model predictions in *Folsomia candida*

**DOI:** 10.1101/2024.08.26.609669

**Authors:** Tom J. M. Van Dooren, Patsy Haccou, Gerard Hermus, Thomas Tully

## Abstract

Population management requires predictions of extinction risk based on a general understanding of these risks and on system-specific modelling. Life tables, available for numerous populations and species, permit calculating population growth and the construction of multi-type branching process models which predict population survivorship and ultimate extinction probabilities. We exemplify this approach and tailor it to an experimental model to predict extinction probabilities per unit of time.

In age-structured populations, founders from different age classes lead to different predicted extinction probabilities. Age effects interact with environmental effects such as culling levels, which influence population growth rates. We assess the accuracy of predictions based on an age-structured matrix model, in an extinction experiment over an eight-week period on the springtail *Folsomia candida*, with crossed founder age and culling level treatments.

Using parameter estimates from an accessory experiment, the fit of model predictions to observed extinction probabilities was generally good. A modified branching process model which allowed culling events between and at observations reduced prediction error. However, additionally maximizing the likelihood of observed extinction probabilities based on survival and fecundity parameters, or on a parameter which concentrated fecundity within a subinterval, did not significantly reduce prediction error according to the AICc.

Our study shows that satisfactory predictions of establishment probabilities and of the initial persistence of small populations can be made using multi-type branching processes and available parameter estimates. Predictions can be improved by integrating knowledge of when events occur within intervals. This can be done without additional parameter estimation.

## Introduction

There is much interest in predicting the establishment or extinction of initially small populations (Stacey & Taper 1992, Thomas et al. 2004, Collen et al. 2011, Kimmel et al. 2022). The relevance is obvious for epidemiology (Jacob 2010, Diekmann et al. 2013), but it is also important in conservation biology and for population management in general (Beissenger & McCullough 2002, Mills 2012). Predictions can be qualitative, for example, whether a population can establish from an initially small number of individuals with a positive probability or not. Quantitative predictions can also be made, for example, cumulative extinction probabilities per unit of time. Many published models in ecology are matrix population models (Caswell 2001, Salguero-Gómez et al. 2015, 2016), which are actually transposed mean matrices of multi-type (Bienaymé-Galton-Watson) branching processes (Athreya & Ney 1972, Caswell 2001, Haccou et al. 2005). For populations where a model with given or adjusted parameter values applies, the population growth rates of such mean matrices directly inform whether ultimate extinction probabilities are one for populations started from a few individuals (growth rate below one, sub-criticality) or smaller (growth rate above one, super-criticality, Athreya & Ney 1972). Probabilities of extinction per time interval of small populations can be predicted using one or several multi-type branching processes corresponding to a given matrix model (Caswell 2001) and this extends to Integral Projection Models (Schreiber & Ross 2016). Multi-type branching processes reduce the need for individual-based simulations (such as in Legendre & Clobert 1995, Ferrière et al. 1996) to make predictions for small populations (Gosselin & Lebreton 2000, Caswell 2001, Schreiber & Ross 2016). How well such processes built upon a mean matrix predict extinctions has to our knowledge not been investigated.

Studies on factors affecting extinction probabilities using replicated populations and associated assessments of predictions are not numerous. As multiple introductions in the wild (Grevstad 1999) are often not feasible, theory coupled to laboratory experiments has provided much of our understanding of demographic, environmental and genetic factors affecting extinction risks in small populations (Griffen & Drake 2008). Several potential determinants of establishment success of small populations have already been experimentally investigated. These include population properties such as initial population size (Belovsky et al. 1999, Drake et al. 2011), between-generation mortality variation (Vucetich et al. 2000), whether individuals are introduced simultaneously or sequentially in an environment (Pike et al. 2004), the immigration rate in small populations (Drake et al. 2005), or for example interspecific predation (reviewed by Griffen & Drake 2008).

Life history traits, and especially survival and fecundity schedules, are major determinants of population growth (Caswell 2001). Differences in population composition with respect to life history traits also affect the risk of extinction (Goodman 1967). Using individual-based simulations of a matrix population model applied to the life cycle of the griffon vulture, Sarrazin & Legendre (2000) demonstrated that introductions of adults were more efficient than juveniles. Even while populations initiated with individuals in different states have the same population growth rate when the population structure has stabilized, the age, size or stage of individuals composing a very small introduced population should be crucial in determining its initial risk of extinction or eventual establishment success.

Comparative studies based on field population data and environmental data aim to identify risk factors for extinction in a correlative manner (e.g. Schwartz et al. 2006, Collen et al. 2011, Saatkamp et al. 2018), but these analyses are not necessarily using life history data via a structured population dynamical model. Jeppsson & Forslund (2012) implemented multi-type branching processes to investigate effects of life history differences in simulations of extinction probabilities, controlling population growth rate by adjusting juvenile survival. Gosselin & Lebreton (2000) applied an age-structured branching process approach to model population numbers in a small population of white stork and Lebreton et al. (2007) simulated extinction probabilities for the Amsterdam albatross. These studies show the potential of an approach where multi-type branching processes are built upon a matrix model. Wider application of this approach, for example for population management, would benefit from an assessment of the reliability of the predictions it makes.

Here, we combine a muti-type branching process approach with an experimental investigation of extinction probabilities in replicated populations each initiated from a single clonally reproducing individual. As an example of the construction and assessment of predictions of extinction based on matrix population models, we use the parthenogenetic model springtail *Folsomia candida* (Willem 1902) to parameterize a matrix model and an age-structured multi-type branching process model based on it. We added a management action to this model, namely culling, and used it to generate predictions of extinction probabilities. These were then compared with observations of population extinctions in an experiment with four levels of culling treatments crossed with three age classes of the single initial individual founding each population. The model made adequate predictions for this experimental system. Using a modified branching process more tailored to the population control effectuated, improved predictions. This suggests that short-term predictions of extinction probabilities based on a parameterized age-structured matrix model can be sufficiently reliable and that the approach deserves to be probed and used more widely in a range of systems and management schemes.

## Material and methods

We first introduce the organism of which we describe the population dynamics by a matrix population model. The parameterization of this model is presented and how parameter values and model structure were translated into a multi-type branching process model. We re-analysed data from (Tully 2023) to obtain parameter values for the modelling. We then describe the experiment used to test predictions of this multi-type branching process models and a slightly different modified model. We present the statistical analysis of the experiment and comparisons between predictions of different models. For analysis, R (R Core Team 2021) was used throughout.

### The springtail *Folsomia candida*

We used the parthenogenetic springtail *Folsomia candida* (Collembola, Isotomidae, Willem 1902) as an experimental model organism (Fountain & Hopkin 2005). Different clonal strains of this springtail have been shown to belong to two main clades (Tully et al. 2006, Tully & Potapov 2015) with different life history traits (Stam et al. 1996, Tully & Ferrière 2008, Mallard et al. 2015, Tully 2023). Within each of the two clades, the genetic, morphological and life-history differences between lineages are negligible (Tully et al. 2006, Tully & Potapov 2015, Tully 2023).

The springtails were maintained in standard rearing containers made of a polyethylene vial (diameter 52 mm, height 65 mm) filled with a 30 mm layer of plaster of Paris mixed with 600µL of Pebeo® graphic Chinese ink. The plaster was kept moist to maintain about 100% relative humidity in the containers. The ink served to increase contrast and to enhance the visibility of the white individuals and batches of eggs. All individuals were kept at 21°C and food (pellets of dried yeast, see Tully & Ferrière (2008) for details) was provided and replaced regularly to ensure *ad libitum* food provisioning.

### Age-Structured Matrix Model

We used individual life history data to parameterize an age-structured matrix model ***A*** (Eqn. 1, Leslie 1945, Caswell 2001) with a fixed interval of one week between successive censuses. On the basis of observed age differences in demographic parameters relevant to the experiment detailed below and considering the duration of the experiment, we chose to use six age classes. We did not aim to incorporate ageing effects at much older ages than six weeks (Tully 2023) because we were interested in what happens in early phases of population establishment starting from young immature or relatively young mature individuals.

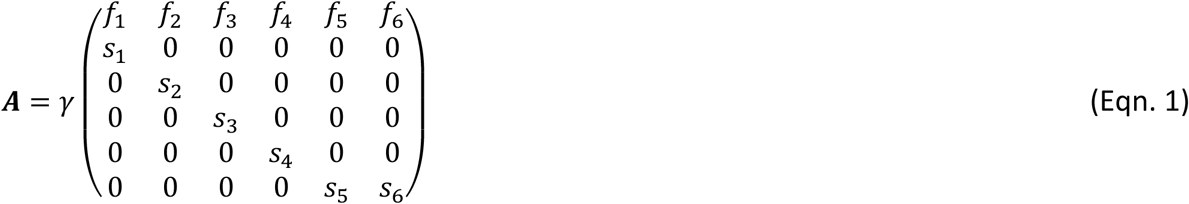

Per age class *j*, the model contains a survival parameter *s*_j_ and a fecundity parameter *f*_j_. We allowed individuals to remain in the oldest age class. This assumption requires that fecundity and survival stay relatively constant from then on, at least for durations relevant to the experiment. The model (Eqn. 1) also contains a culling parameter **γ**, the probability that an individual or clutch is surviving a culling event applied to the entire population right before the census.

### Estimation of Survival and Fecundity Parameters

We reanalysed life history data collected in five closely related strains (labelled “GM”, “PB”, “TO”, “US”, “WI”, Tully 2023). Ten individuals per clone were isolated just after hatching and kept in the previously described individual rearing vials and fed ad libitum. Repeated observations allowed us to determine individual lifespans, the times of clutch laying, the numbers of eggs and newborns per clutch. In this experiment, durations of egg development were not determined. For calculations below, we used the median duration of egg development reported in (Tully 2004) to be 9.5 days for the GM (“Grotte de Moulis”) clone, which is the clone used for the extinction experiment detailed below.

From the individual life history data, age-class fecundities *f*_j_ and survival probabilities *s*_j_ (Eqn. 1) were estimated for the GM clone on the basis of random effects models fitted to the data on all five clones.

To obtain age-specific survival probabilities *s*_j_, we first estimated the baseline hazard per hour using the five clones and modified it with a random effect for the clone GM (Supplementary Material). The baseline hazard was estimated using a kernel smoothed method (Muller & Wang, 1994, Supplementary Fig. S1). Using the median development time per egg, we can assume that an individual has age zero when it is laid as an egg, age 9.5 days when it hatches and that the individual survival and fecundity data describe patterns starting at age 9.5 days. Probabilities of survival before hatching for the clone GM were estimated based on the fraction of eggs which did not hatch. Given the low mortality of recently hatched individuals (Fig. S1), we assumed that death before hatching occurred within the first week of age and with a constant rate. To obtain six *s*_j_ values from probabilities *l*(j) of survival up to age *j* in weeks, we used approximations for birth-flow populations (Caswell 2001, supplementary material)

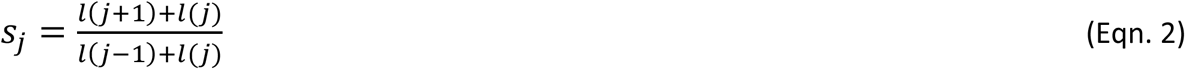

For fecundities *f*_j_, we first fitted the clutch sizes per hour per individual as a function of age after hatching with zero-inflated mixed generalized linear Poisson models (*mgcv* library Wood et al. 2016). These yielded an age-specific and clone-specific probability of producing a clutch per hour and an age-specific clutch size, conditional on producing a clutch. In order to obtain fecundities per week for each age class, we predicted numbers of eggs produced per hour using model predictions for clone GM, for ages occurring within this experiment. We set numbers of eggs to zero for ages below which no reproduction had been observed in any of the clones (age 3.14 weeks). We then used an approximation for birth-flow populations (Caswell 2001, Supplementary Material) applied to the predicted response.

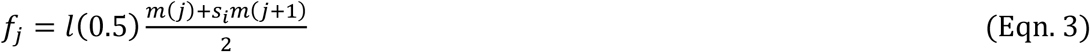

where *s*_*j*_ is the survival probability in age class *j* (Eqn. 2) and *m*(*j*) the summed predicted number of eggs produced by a single individual, summing all hourly values from age *j* - 1 until (not including) age *j* in weeks. Note that in this parameterization (Eqn. 3), fecundity parameters are not independent of survival of the parent.

### Branching Process Modelling

Once the matrix model ***A*** parameterized, we investigated its eigenvectors and eigenvalues to determine population growth rates, reproductive values and stable age distributions for different values of the probability to survive culling **γ** (Supplementary Material). The dominant eigenvalue **γ**(***A***) of the matrix model corresponds to the long-run population growth rate per week. According to branching process theory (Athreya & Ney 1972), a population with individual variation in fecundity for which the mean matrix is the transpose of an age-structured matrix model will certainly go extinct in the long run if the largest eigenvalue is less than or equal to one (**γ**(***A***) ≤ 1, subcritical or critical cases). If this eigenvalue is larger than one, extinction occurs with a probability less than one (**γ**(***A***) > 1 supercritical cases). An initially small founder population, even of a single individual, then has a chance to escape from extinction and establish itself. In the first case, we can expect survivorship curves to decrease to zero survivorship. In the second case, a horizontal asymptote will occur at a non-zero survival probability and a fraction of populations will become established, i.e., without further risk of extinction when environmental change or population regulation don’t reduce survival nor fecundity.

We wanted to predict not just the possibility to escape extinction, but also the probability of extinction per unit of time. Therefore we developed a multi-type branching process model (Athreya & Ney 1972, Supplementary Information) based on the age-structured matrix model (Eqn. 1), where we assume that (1) individuals in age class *j* survive between censuses according to independent Bernoulli distributions with mean *γs*_*j*_, (2) clutches survive between censuses according to independent Bernoulli distributions with mean *γ*, and (3) that newborns hatch from clutches produced by age class *j* in numbers specified by Poisson probability distributions with mean *f*_j_. Assuming in addition that survival and reproduction between censuses are independent, this leads to a simple multi-type branching process model (Caswell 2001, Lebreton et al. 2007, Jeppson & Forslund 2012).

Probabilities of extinction or of establishment are calculated using generating functions, which, depending on what is calculated from them, are also called probability generating functions or reproduction generating functions. Probability generating functions permit efficient calculations of probabilities of events and the study of properties of the probability distribution functions of discrete random variables. Here, the multi-type branching process model is specified by a multivariate (or vector-valued) generating function ***g***(***x***) = (*g*_1_(***x***), *g*_2_(***x***), …)where ***x*** is a vector with six elements. Each *g*_*j*_(***x***) = *E*[***x***^***z****j*^] is the generating function for age class *j*, with vector ***z***_*j*_ = (*z*_1j_, *z*_2j_,…, *z*_6j_) in the exponent containing the numbers *z*_ij_ of type *i* individuals produced by a single type *j* individual across one time interval and E[] denoting taking the expectation with respect to the probability distribution function of ***z***_*j*_ (Supplementary Material). From this multivariate function, predicted numbers of individuals present per different age classes at the next census can be calculated, starting from a population with a single individual in one age class or starting from populations with individuals in different age classes. Given the assumptions above, the Supplementary Material shows that for *j* in 1, …, 5,

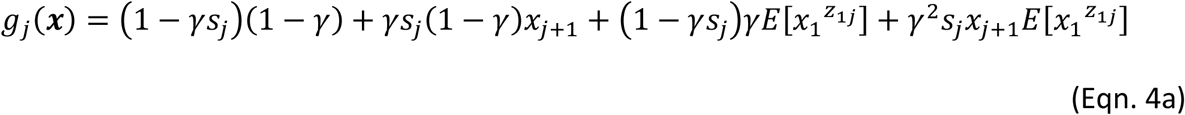

and for the oldest age class *j* = 6,

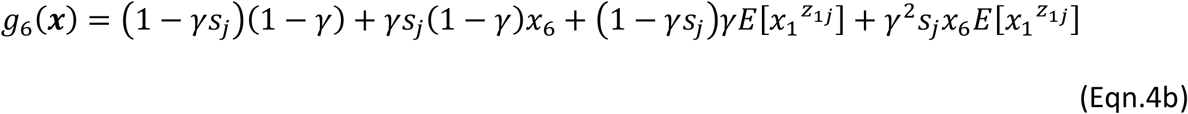

Here we note that 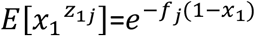 (Supplementary Material).

This probability generation function depends on ***x*** and on parameters *s*_j_, *f*_j_ and **γ**. For a population initiated from a single individual of type *j*, we denote the numbers of individuals in different states *i* in a population at time *t* (*t* a nonnegative integer) by a vector ***Z***_*j*_(*t*), with elements *Z*_*ij*_(*t*) (*i* = 1,…, 6). The corresponding vector-valued probability generating function ***G***(***x***,*t*)contains entries 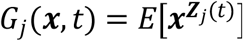. A useful recursion relates these to ***g***(***x***) (Supplementary Material), which is

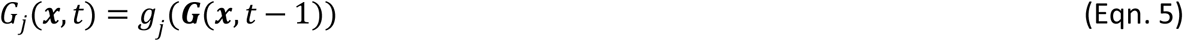

### Subdivision of Time Intervals

When culling does not happen right before census but somewhere at a fixed time within the interval between observations, we don’t expect this to change the population growth rate, because the underlying process is unaffected and only the moments of observation change. Therefore, whenever culling happens within an interval, as long as it is always at the same duration after observation, the model parameter values distinguishing super-from subcriticality should remain the same but probabilities of extinction per interval not. Culling now divides an interval of length *t* between observations in two intervals of lengths *t*_1_ and *t*_2_ = *t* - *t*_1_. Probability generating functions can be defined for each separate interval, which together compose the probability generating function across the entire interval ***h***(***x***) = ***h***_**1**_(***h***_**2**_(***x***))(Supplementary Material). To arrive at a branching process model integrating processes in the two subintervals and also slightly extending the model (Eqn. 1, Eqn. 4), it is now assumed that age class one produces no clutches. This allows us to work out the generating functions ***h***, with ***x*** and ***z*** which are ten-tuples. Developing embryos at the moments of observation are now modelled as clutches (*z*_1_ to *z*_5_ are the numbers of clutches produced by age classes two to six), and the numbers of hatched individuals (five additional states with hatched individuals of age classes two to six *z*_6_ to *z*_10_). Clutches can be produced by all individuals present at the start of subintervals. An individual in state *j*, corresponding to age class *j* – 4, produces a clutch within *t*_1_ with probability *p*_j-4_, if not, it will do so in *t*_2_ when still alive at the start of that interval. Probabilities *p* are equal to zero for age classes with zero fecundity. It is assumed that clutches and individuals can be culled at the end of each subinterval, with probabilities *γ*_1_and *γ*_2_. Survival of clutches is assumed to be equal to one in the absence of culling. Of individuals in state *j* corresponding to age class *j* - 4 it is *s*_1,*j*−4_ and *s*_2,*j*−4_ in the two subintervals respectively, where it is assumed that *s*_1,*j*_*s*_2,*j*_ =*s*_*j*_. Culling of an entire clutch already present at the start of an interval can occur if the eggs don’t hatch within the interval. A clutch produced by an individual of age class *j* - 4 in the previous time interval survives until after the first culling within the interval with probability *γ*_1_. It is assumed that all eggs from such a clutch hatch within the second subinterval and recruit to the second age class (which is the sixth state here). We assume that the number of hatched individuals surviving until after the second culling is Poisson distributed with mean *s*_1_*f*_*j*+1_*γ*_2_. In the Supplementary Material, the generating functions *h*_*j*_(***x***)are given and one can derive that the matrix population model corresponding to the new assumptions is

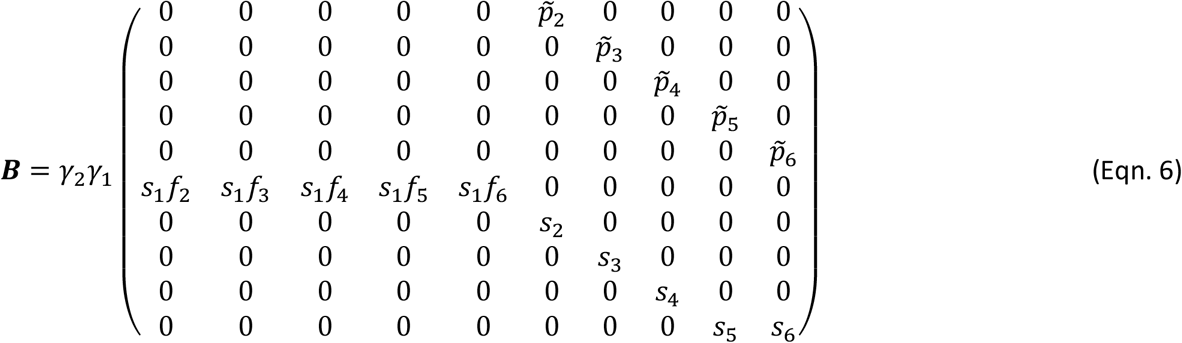

With 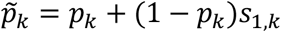, the probability to produce a clutch within an entire interval. The vector of numbers projected forward by repeatedly applying ***B*** consists of the expected numbers of clutches produced by each age class (two to six) and the expected numbers of hatched individuals in age classes two to six. In total these are numbers of individuals in ten individual states.

Assuming that *γ*_2_ and all *p* are equal to one and that *γ*_1_ =*γ*, one can find that expected lifetime reproductive success of a hatched individual observed in state six is 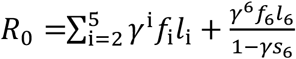. This is the same expression as the expected lifetime reproductive success corresponding to (Eqn. 1), when assuming that *f*_1_ is equal to zero. Expected lifetime reproductive success can be used equivalently to population growth rate to predict super- and subcriticality (Jagers & Sagitov 2000, Supplementary Material). Therefore, when culling occurs after a subinterval and not at the end of an entire interval and if one assumes that reproduction and survival are independent across the entire interval, models ***B*** (Eqn.6) and ***A*** (Eqn. 1) make identical predictions in this regard.

### Calculation of Survivorship Functions

We determined survivorship functions, i.e. the probability that the population is not extinct at each census, for populations founded by a single individual (Supplementary Material).

These survivorship functions depend on the age class of the founding individual and on the demographic parameters. We briefly summarize how to obtain survivorship functions (Athreya & Karlin 1971, Caswell 2001). The survivorship function for initial condition *j*, i.e., a population founded by one individual in state *j* at time zero is defined as:

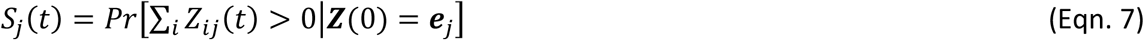

with ***Z***(0)the population state vector at time zero. We can determine *S*_*j*_(*t*)as follows for a given multi-type branching process, ***g***(***x***) is used as an example here. Let *Q*_j_(*t*) denote the probability of extinction by time *t* of a population founded by a single type-*j* individual and ***Q***(*t*) = (*Q*_1_(*t*), …, *Q*_6_(*t*))the vector with the six cumulative extinction probabilities *Q*_j_(*t*). The probability *Q*_j_(*t*) turns out to be a simple recursion of the probabilities at time *t* − 1, i.e., *Q*_*j*_(*t*) = *g*_*j*_(***Q***(*t* − 1))(Supplementary Material). ***Q***(0) = **0**, and all further ***Q***(*t*)can be obtained by recursion using the vector-valued probability generating function (Eqns. 3a & 3b). Survivorship is the probability of not being extinct by time *t*. It is therefore equal to *S*_*j*_(*t*) = 1 − *Q*_*j*_(*t*). From a survivorship function, the probabilities that a population goes extinct across one weekly interval ending at time *t* can be calculated for each interval between censuses. They are equal to *p*_*j*_(*t*) = *S*_*j*_(*t*)/*S*_*j*_(*t* − 1).

### Sensitivity Analysis

From a matrix model, sensitivities of growth rates to model parameters can be derived using standard expressions (Caswell 2001). For ***A***, we calculated sensitivities of population growth rate to fecundity and survival parameters and to culling parameter **γ** (Supplementary Material). For different culling levels, it can be shown that these are proportional to culling level and therefore preserve the pattern across fecundity and survival parameters for each culling level (Supplementary Material). We used the branching process model ***g***(***x***) to investigate the sensitivity of extinction probability predictions to model parameters. For each of the demographic parameters in the model, we determined how small changes in them will affect the ultimate extinction probabilities 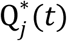 which are the equilibrium values of the discrete-time dynamical system ***Q***(*t*) = ***g***(***Q***(*t* − 1))with initial condition ***Q***(0) = **0**. Extinction probabilities of populations founded with one individual of a single age class were estimated for small parameter increases and small decreases of one to four percent and per culling level in the experiment. For fecundities *f*, we used the results for increases. For survival probabilities *s* and **γ** we used the results for decreases. A linear regression was fitted to the set of changes per parameter and the estimated slope is the sensitivity of ultimate extinction probability to the parameter for a specific age class.

### Two-way Population Establishment Experiment

We established populations founded by a single individual of clone GM to assess how well the multi-type branching process model(s) predict survivorship and extinction probabilities. Predictive capabilities might vary with environmental or population state and to this end we introduced two crossed treatments.

#### Age class of founding individual

We replicated populations, each founded with a single female individual of known age class. We used three founder age classes, which, according to our theoretical predictions, should result in different outcomes in terms of extinction/establishment probabilities. Prior to the start of the experiment, we grew cohorts of the required age classes by isolating eggs laid by stock populations in a one-week interval, by providing food once all eggs had hatched, and by sampling individuals at weekly censuses after isolation of the eggs. The founding individuals were thus of age classes two (newborn), four (small adult), and six (large adult) according the matrix population model (Eqn. 1).

#### Culling

To each population, we applied a weekly culling treatment with three different intensity levels and an unculled control. Culling started four days after the populations were founded. Culling was accomplished by randomly selecting a quarter, one half or three quarters of the surface of the rearing substrate and by removing all eggs and individuals from this selected area (Pike et al. 2004). In the control treatment, no culling was carried out (**γ** = 1). In three other treatments, the proportions culled were either 25%, 50% or 75%, parameterized by the probabilities of surviving culling (**γ** = 0.75, 0.5 and 0.25). Populations were checked for extinction at weekly intervals, each time, three days after culling. Not all of the populations were founded on the same date and different sets were censored at different times. Forty populations were censused eight times (all initiated with stage six individuals), 119 populations until individuals were seven weeks old, 81 five times and 60 four times. For each combination of culling level and founder stage, 25 replicate populations were initiated (300 populations in total). All data are available as Supplementary Material.

### Statistical Analysis Two-way Experiment

We tested for differences between survivorship curves using the scores of Sun (1996). Interval-censored survival analysis (Collett 2014, Bogaerts et al. 2017) was used to compare treatments and test for time-dependent effects. This was done using binomial generalized linear models (GLM), with a complementary log-log link function (Harney et al. 2013) for the probability of extinction per week. In the analysis, we included categorical effects of *founding stage, culling level* and categorical or continuous effects of *census week*.

Likelihood ratio tests were used to simplify the models. The data are Bernoulli presence-absence data, but there is no covariate which is specific to an individual population. Therefore, overdispersion could occur and we assessed this by fitting quasi-binomial models and examining the overdispersion parameter estimate. We found no evidence of overdispersion. Multiple comparisons of estimated parameters in the GLM were performed using simultaneous confidence intervals following Bretz et al. (2001).

We determined for which culling treatments population growth rates were supercritical or subcritical using the matrix model ***A*** (Eqn. 1). For each treatment combination in the experiment, the branching process models ***A*** (Eqn. 1) and ***B*** (Eqn. 6) predict the expected survivorship of a population. We compared these predictions visually with Kaplan-Meier survivorship estimates adjusted for interval-censored data. We compared the deviances of the data with respect to each model.

### Re-estimation of Model Parameters

We fitted the extinction probabilities predicted by the branching process models to the extinction probabilities observed in the experiment using maximum likelihood (ML). We maximized the likelihood of a subset of model parameters given the data, restricted to parameters for which the sensitivity analysis indicated that they affected extinction probability. The optimized predictions are compared with our initial predictions based on the accessory experiment using AICc (Hurvich & Tsai 1989), an information criterion for small sample sizes.

## Results

### Demographic Parameters

We did not observed any reproduction before 299 hours after hatching (12.5 days), therefore the minimum observed age at reproduction since laying was 527 hours (22.0 days). The values of survival and fecundity parameters obtained using birth-flow population approximations (Caswell, 2001) are given in Table 1. In the absence of culling, the population growth rate per week is 3.21 for these parameter values and model ***A*** (Eqn. 1). The branching process is critical for **γ** = 0.31. If we assume that culling occurs as in the experiment, four days after census, and not immediately before census, that reproduction occurs with probability 4/7 after census and before culling, then the population growth rate for model ***B*** (Eqn. 6) is equal to 3.19 and the branching process is critical for **γ** = 0.31 as well. If we additionally assume that mortality within an interval is constant, such that the survival probabilities per day are 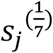, then we can also predict extinction probabilities per week for model ***B*** using a multi-type branching process (Supplementary Material Eqns. S.9). Figure 1 shows that predicted cumulative extinction probabilities are generally larger when processes within the interval are modelled, and that differences between predictions for founders in age classes six and four are small for each model.

**Table 1.** Life history parameters of *Folsomia candida* obtained from data in Tully (2023) and Tully (2004) using birth-flow approximations (Caswell 2001) for survival probabilities and fecundities. Population growth rate sensitivities are added. For survival probabilities and fecundities, the value given assumes **γ** = 1.

|  | Model parameter | Value | Growth rate Sensitivity |
| --- | --- | --- | --- |
| Survival probability per week | $s_1$ | 0.994 | 0.824 |
| | $s_2$ | 0.997 | 0.821 |
| | $s_3$ | 0.977 | 0.571 |
| | $s_4$ | 0.943 | 0.232 |
| | $s_5$ | 0.913 | 0.073 |
| | $s_6$ | 0.887 | 0.027 |
| Fecundity | $f_1$ | 0 | 0.248 |
| | $f_2$ | 0 | 0.077 |
| | $f_3$ | 11.15 | 0.024 |
| | $f_4$ | 43.71 | 0.007 |
| | $f_5$ | 68.54 | 0.002 |
| | $f_6$ | 74.50 | 0.001 |
| Survival probability culling | $\gamma$ | Controlled | 3.206 |
| Hatching | Age in days | 9.50 |  |
| Earliest reproduction | Age in days | 21.96 |  |

**Figure 1.**
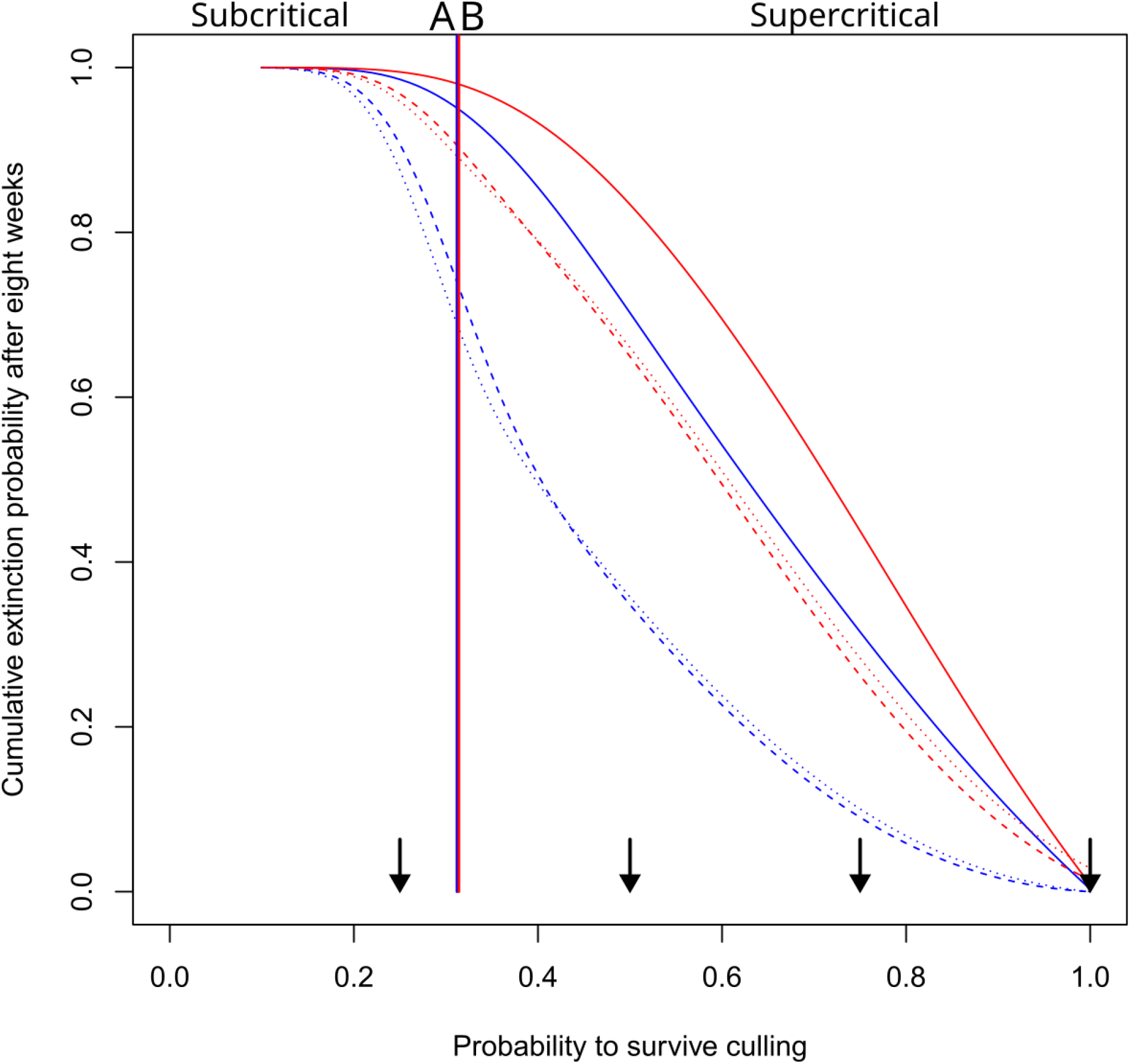
Model predictions of extinction probabilities. Predicted cumulative probabilities of extinction eight weeks after founding are shown as a function of the probability to survive a culling event. Eight weeks is the longest duration populations were observed in the extinction experiment. Results are shown for the branching process models based on model **A** (Eqn. 1, blue) or the branching process leading to matrix ***B*** (Eqn. 6, red), when founders are either in age class two (full line), age class four (stippled line), age class six (dotted line). Vertical lines indicate the value where the branching process is critical. Arrows denote culling intensities in the extinction experiment.

### Sensitivities

The population growth rate is most sensitive to survival in the first age classes (Table 1). Small changes in the fecundities in age classes four to six have negligible effects on population growth (Table 1) and ultimate extinction probabilities (Figure 2). For ultimate extinction probabilities, sensitivities to survival and fecundity and to the probability to survive culling are generally negative (Fig. 2), so any increase in these parameters reduces the probability of ultimate extinction. For supercritical parameter combinations, the extinction probability for a population initiated with an individual in age class *j* is most sensitive to the survival probability *s*_j_ (Figure 2). A similar pattern occurs for fecundities, where the extinction probability for populations founded by age class *j* is most sensitive to *f*_j_. Fecundities have overall smaller values of the sensitivities.

**Figure 2.**
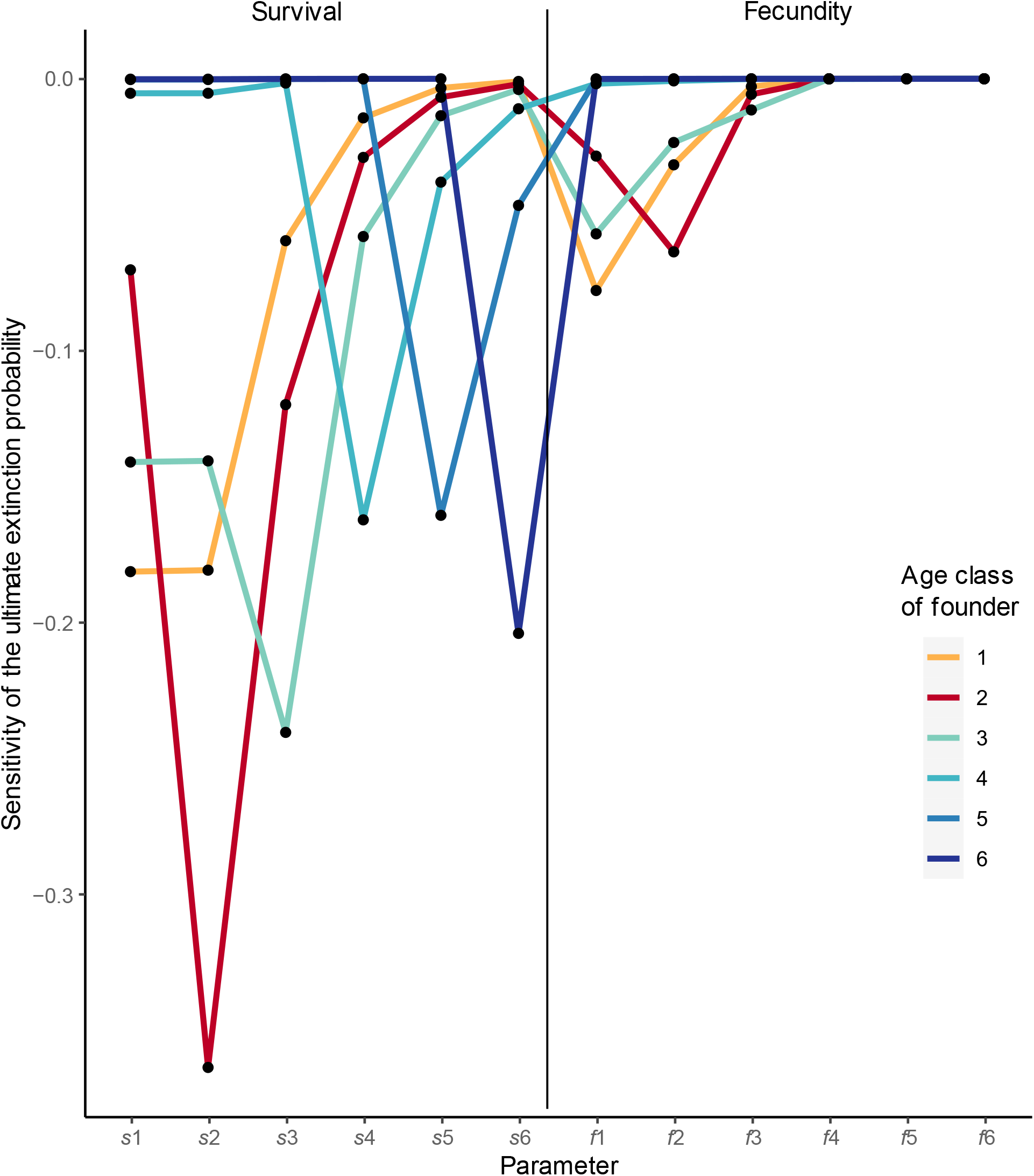
Sensitivities of ultimate extinction probabilities. Sensitivities of the ultimate extinction probabilities to the different survival parameters (*s*) and to fecundity parameters (*f*). The sensitivities have been estimated for the branching process model based on ***A*** (Eqn. 1), for a 50% culling intensity (**γ** = 0.5), and for different age classes of the founder. Other parameters are as in Table 1.

### Statistical Analysis of the Extinction Experiment

The control and two of the culling treatments correspond to supercritical regimes, it is only when 75% of a population is culled each week that the corresponding branching process is subcritical (Fig. 1). The survivorship curves in the different treatment combinations are not equal (χ^2^_11_ = 161.7, *p*-value < 0.0001, Figure 3) and when we inspect the survival probabilities after eight weeks, the pattern seems to correspond globally to the expectations based on extinction probabilities in Fig. 1. When the extinction probabilities per week were modelled with a binomial GLM, the selected model had additive effects of week (linear effect χ^2^_1_ = 38.6, p-value < 0.0001), founder age class (χ^2^_2_ = 20.9, p-value < 0.0001) and culling level (χ^2^_3_ = 187.5, p-value < 0.0001). Parameter estimates are given in Table 2. The negative week effect demonstrates that extinction probabilities decrease over the course of the experiment (Table 2). When we tested which founder age classes and culling levels had different extinction probabilities using post-hoc pairwise comparisons, we found that all culling levels had significantly different effects on extinction probability. As expected (Fig. 1), the extinction probability increases with culling (Table 2). The extinction probability was significantly lower for populations founded by age class four than by age class two (difference between parameter estimates −0.918 (s.e. 0.217), *p* < 0.0001), and lower in age class six than in age class two (difference between parameter estimates −0.728 (s.e. 0.212), *p* = 0.0016). Extinction probabilities did not differ between founder age classes four and six.

**Table 2.**
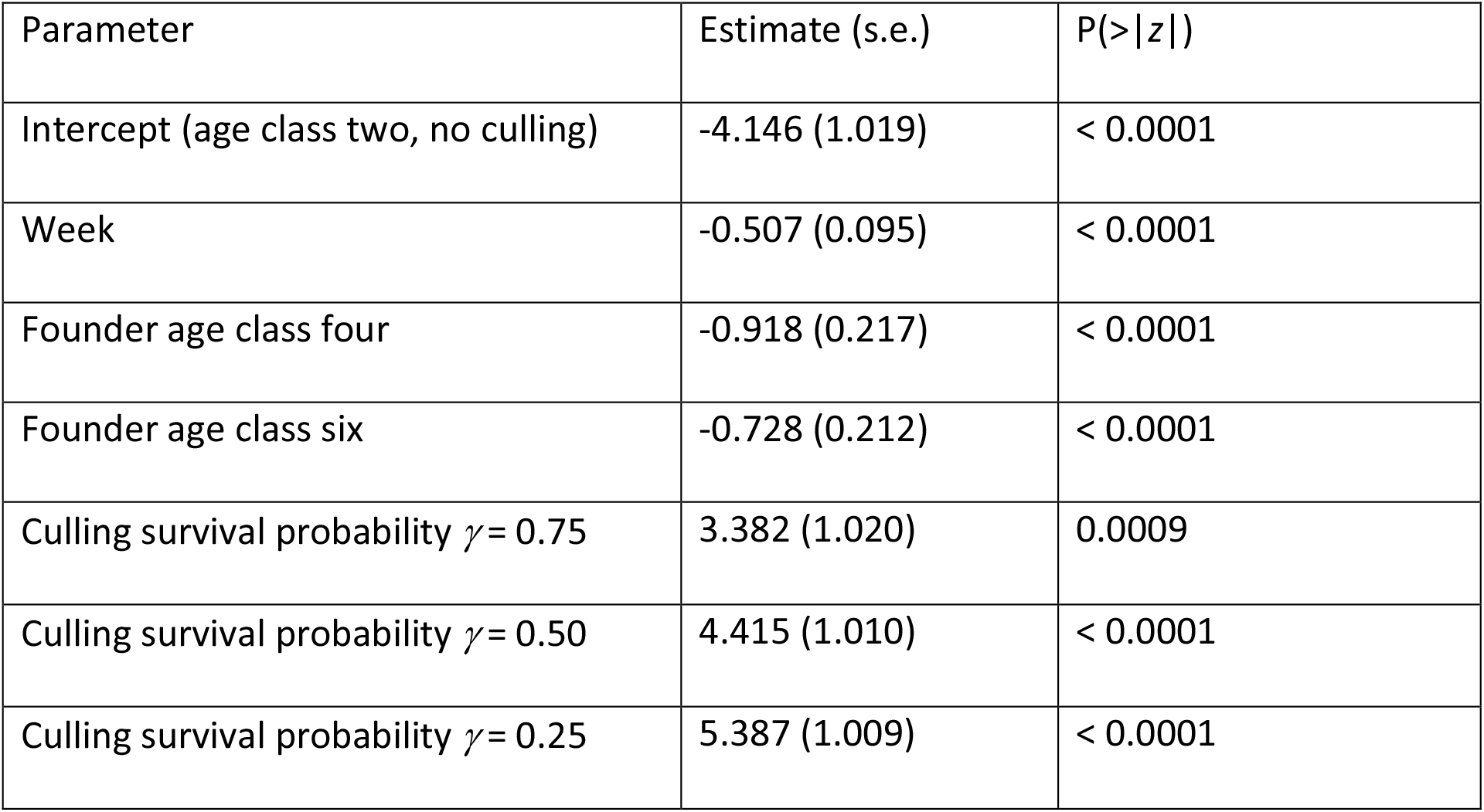
Parameter estimates of the selected GLM. An overview of parameter estimates and their standard errors of the binomial generalized linear model GLM for extinction probabilities. This model was obtained after model selection. An extra column indicates which parameters are significantly different from zero according a *z*-test.

**Figure 3.**
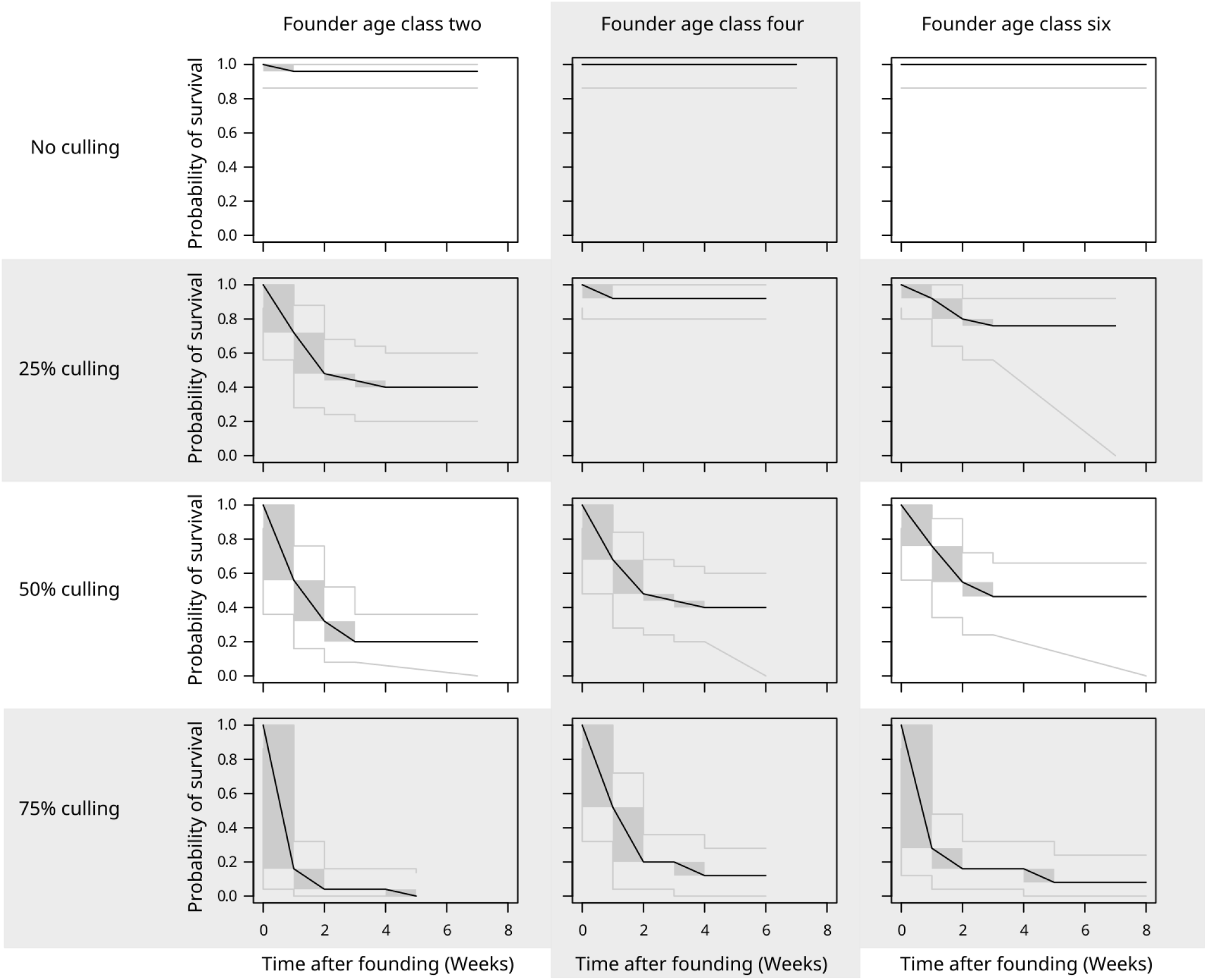
Survivorship curves for different founder and culling treatments. Fractions of non-extinct populations as a function of time since the beginning of the experiment, in weeks. The data are interval-censored and the plotted curves are therefore indeterminate for some time intervals (Bogaerts et al. 2007). They are then drawn as boxes with a diagonal line. Nonparametric maximum likelihood estimates NMLE of the survivorship curves are plotted as black lines. Intervals where the nonparametric maximum likelihood estimate is indeterminate are drawn as gray rectangles, and for readability, a black line is drawn from the upper left to lower right corner of each rectangle. Confidence intervals for the survivorship curves are plotted as grey lines.

### Survivorship Predictions

When using the parameter estimates (Table 1) to predict survivorship curves, and assuming that culling occurs immediately before the census, we found that our predicted survivorship curves stayed within the confidence band of the non-parametric survivorship estimates for ten out of twelve treatment combinations for model ***A*** (Figure 4) and for eleven for model ***B*** which models the timing of culling more precisely. The deviance of model ***A*** equals 532.6 and of model ***B*** 496.7. No parameters are estimated within the experiment, so we can interpret these as AICc. When we optimized the likelihood of the observed extinction probabilities as a function of all six survival parameters and fecundities in age classes two and three, we found that the AICc became 519.9 and 507.9, respectively. Optimizing the likelihood just in function of the probability of producing a clutch within the first subinterval gives an AICc of 498.3. None of these information criteria are below the deviance of model ***B*** without optimization, therefore the a priori constructed model which takes subintervals into account is preferred and is expected to predict the data better.

**Figure 4.**
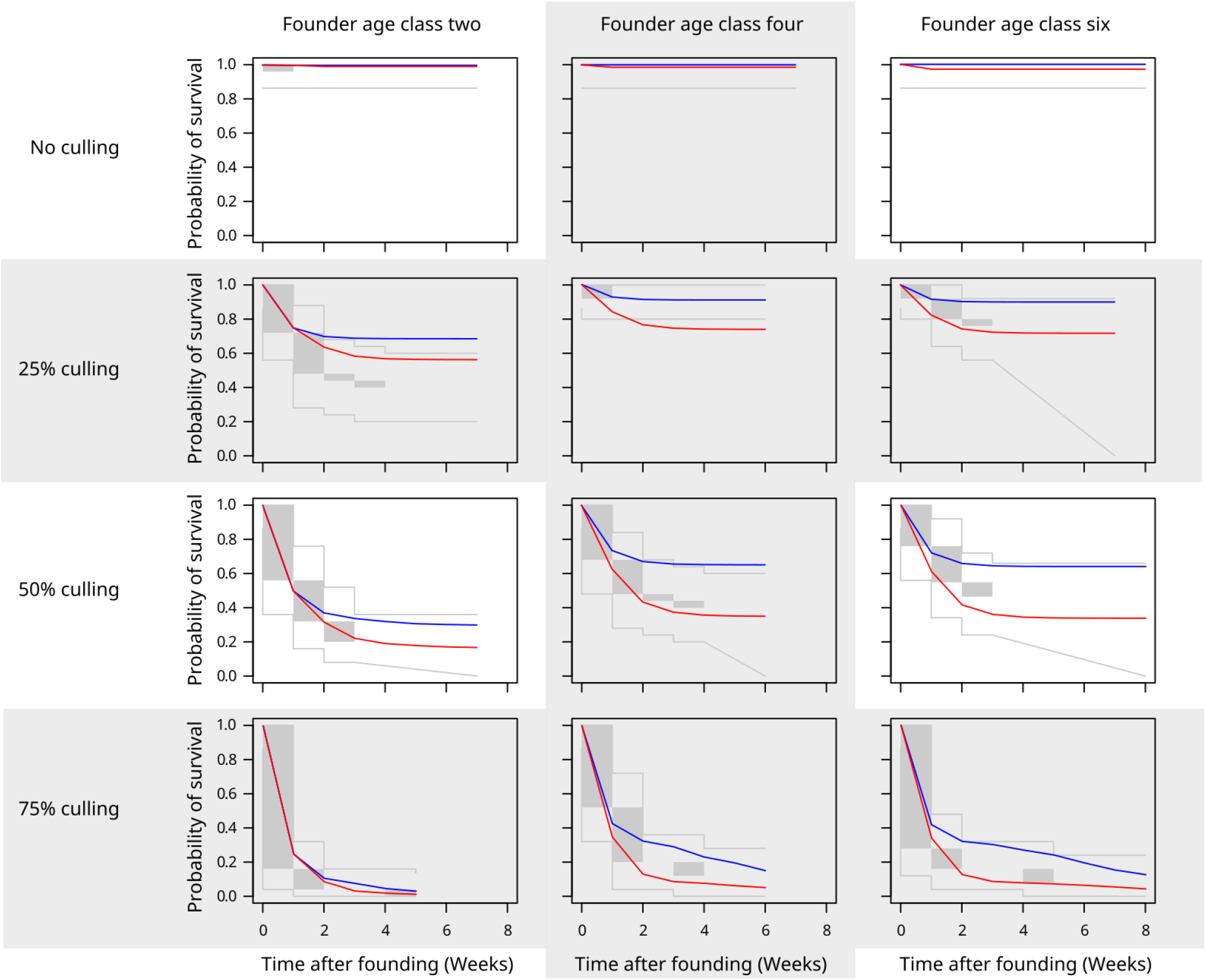
Comparisons of model predictions with survivorship. Confidence intervals of survivorship curves (grey) are shown for different treatment combinations in the culling experiment (i.e., Fig. 3 with NMLE removed). Predictions of the branching processes based on model ***A*** (blue) and model ***B*** (red) are superimposed for each treatment combination. The treatments in the lowest row correspond to subcritical cases (Fig. 1).

## Discussion

Matrix population models (Caswell 2001) are essential tools in population ecology. It is less commonly known that they are actually transposed mean matrices of multiple multi-type branching processes (Athreya & Ney 1972, Caswell 2001). We derived such a branching process from a parameterized matrix model and used it to predict extinction probabilities in an experiment with replicated small populations each founded by a single clonal individual, to which we applied twelve treatment combinations. The predictions were satisfactory but could be improved by changing the branching process model so that it accommodated the fact that reproduction and survival within an interval were not necessarily independent, due to culling occurring about halfway each interval. Re-estimating model parameters did not reduce the AICc values, therefore such adjustments did not significantly improve the fit of the model to the data and its predictive performance.

### Matrix Population Models

Given a branching process model, a first assessment of extinction probabilities occurs by determining whether its mean matrix has a supercritical growth rate or not. In a similar manner, each matrix population dynamical model can be used to determine for all corresponding branching processes when they would be supercritical, critical or subcritical. We used the expected lifetime reproductive success *R*_0_ to determine supercriticality (Jagers & Sagitov 2000) and found that for the more detailed branching process with matrix ***B***, the same expression for *R*_0_ could be recovered, if we assume that reproduction and survival are independent in each entire interval and that culling occurs only once within an interval. Without estimating new parameters, i.e., by assuming constant mortality rates within an interval and assuming that fecundity (the production of a clutch) in a first or second subinterval is proportional to their duration, we observed that *R*_0_ values for models ***A*** and ***B*** became equal to one at very similar culling survival probabilities **γ** (Fig. 1). Different proxies of population growth rate such as *R*_0_ might be of much use to explore for a set of models at which parameter values their growth rates are critical. However, for instantaneous extinction probabilities from week to week, models ***A*** and ***B*** did differ (Fig. 1) and branching process predictions are required to predict the fate of populations in the short term.

### Culling Intensity

As expected, increasing levels of culling modified population survivorship curves. Culling 75% of a population each week led to a subcritical growth rate and drove populations towards a certain extinction. This treatment can be compared to treatment 6b in Pike et al. (2004), where one third of populations underwent a 25% culling every two days, which is roughly equivalent to our culling treatment of 75% per week. Pike et al. (2004) found that populations under these conditions remained at risk of extinction regardless of their age since initiation, but did not estimate a population growth rate. Model ***B*** predicted more extinctions in the presence of culling, as it occurred earlier within each interval when some individuals had not yet reproduced. We also assumed for model ***B*** that entire clutches could be culled twice, and not just within the week in which they were produced.

### Founder Age Class

The age class of a founder does not affect sub- or supercriticality in the long run. For each treatment with a supercritical growth rate, the predicted extinction probabilities per week of populations founded by different age classes converged to zero at the end of the extinction experiment, at different cumulative extinction probabilities. Note also that for culling levels corresponding to supercritical growth, survival probabilities do differ between populations founded by individuals in different age classes (Fig. 3, Table 2). Starting a population with an adult (age class 4 or 6) rather than a juvenile (age class 2) improves establishment success by reducing the period during which the founder is alone in the population, with no descendants present. This result is similar to that shown for a model applied to griffon vultures (Sarrazin & Legendre 2000, Robert et al. 2004). For populations with founders in non-reproductive age classes, an assumption of independence of survival and reproduction within each interval has no effect, and model predictions of different branching processes assuming this independence or not can only differ later when reproductive individuals are present.

### Heterogeneous Demographic Rates

A branching process model was proposed, model ***B***, which divided a time interval into two subintervals, each of which could have a culling event at the end. In the Supplementary Material it is discussed that such subdivisions can be modelled starting from composite generating functions, with different functions for each of the subintervals. This approach can be extended to model multi-type branching processes with different events and rates per time step, and in general, composite generating functions could be used to model population dynamics in sequences of changing environments (Athreya & Karlin 1971, Dolgopyat et al. 2018), which involve fewer assumptions than the random environments often considered (Athreya & Karlin 1971, Vatutin & Dyakonova 2000). Even without environmental variation, demographic rates in structured population models generally vary between individual states, and it seems worthwhile to further explore the performance of multi-type branching processes derived from matrix models for types of structuring other than age, such as IPM with for example size structure (Schreiber & Ross 2016). In this springtail example, we were able to start populations in the experiment from a single individual because reproduction occurs clonally. Heterogeneity in rates between the sexes should be considered in populations where it applies, and two-sex multi-type branching process models (Mode 1972, Hull 2003, Fritsch et al. in press) might need to be assessed for all systems where matrix population models assume so-called female dominance, which is a common assumption (Lebreton et al. 2007, Salguero-Gómez et al. 2016). Since parameterized matrix models or Integral Projection Models are available for many species, broad comparative assessments of extinction risk of small populations in the short term could make use of appropriate multi-type branching processes and wouldn’t just regress measures of extinction risk on life history statistics.

### Population regulation

For culling treatments corresponding to supercritical population growth rates, extinction probabilities after the first four weeks were small. Our branching process model predicted hardly any extinction thereafter, suggesting that the ultimate cumulative extinction probability could be reached quickly, after which populations could be considered established. These populations are expected to continue to grow exponentially (Athreya & Ney 1972), which is unrealistic in the long term. It is expected that their sizes remain bounded because of population regulation and that demographic stochasticity occurs at any finite population size (Lebreton et al. 2007). Gosselin & Lebreton (2000) elaborated the notion of quasi-stationarity for populations which have a subcritical growth rate or are bounded by population regulation. Conditional on non-extinction and for population regulation which would make a corresponding deterministic model converge to a stable equilibrium (e.g., Block & Allen 2000), numbers of individuals in different states then converge to a quasi-stationary distribution and the extinction probability per unit of time to a constant. We did not explore these issues here in the experiment, as we focused on short-term extinction probabilities and ignored population regulation, which would change model parameter values over time. When the branching process predicted zero extinctions or when we did not observe further extinctions, populations might or might not have reached their quasi-stationary distributions. Whenever a population reaches a state with few individuals where population regulation is negligible again, the multi-type branching process should predict extinction probabilities well for a while.

### Population Management

Concerns over Population Viability Analysis PVA (Beissenger & McCullough 2002), a method used in conservation biology to predict the persistence of populations, have involved overconfidence in modelling results (Reed et al. 2002), even though retrospective tests have shown, for example, the ability of the approach to predict population decline in an unbiased manner (Brook et al. 2000, Ellner et al. 2002). We have compared results of extinction probabilities per time step between two models relevant to the experimental system. The encouraging fit of the predictions by both models to the experimental data calls for a wider use of branching processes derived from matrix models to predict the fate of small populations and to manage them. In the experiment, we managed populations by culling them, and such interventions can be easily incorporated into population models without the need to estimate additional parameters. Other types of management intervention may be relatively easy to add to a model. The comparison of the performance of two branching process models we used suggests that there is some robustness to model misspecification, but applications of the approach to making predictions for a single specific population should take into account the uncertainty in model parameters. For systems with sexual reproduction, care should be taken to model different marriage functions (Caswell 2001) or when not, the assumptions involved should be assessed. Nevertheless, predictions of extinction probabilities before populations grow large, should be based on multi-type branching processes, not on population growth rates and they should therefore be an integral part of population viability analysis, just as results based on quasi-stationary distributions for established populations (Lebreton et al. 2007) should.

## Author Contributions

TT, PH and TVD designed the experiment. PH, GH and TVD developed the model. GH and TVD carried out the extinction experiment. TVD, TT and GH analysed the data. TVD and TT wrote a first draft of the manuscript. All authors commented on subsequent versions of the draft.

## Acknowledgments

TT received support from ANR Evorange (ANR-09-PEXT-011). Souleyman Bakker commented on a previous version of the manuscript.

## Supplementary Information

With: Extinction probabilities of small structured populations: adequate short-term model predictions in *Folsomia candida*. Tom J. M. Van Dooren, Patsy Haccou, Gerard Hermus, Thomas Tully.

### Matrix population dynamical model

In branching process modelling, the mean matrix of a process can be derived from the generating functions (Athreya & Ney 1972). In the other direction, any discrete-time matrix population dynamical model can be used as a starting point to derive a corresponding multi-type branching process in discrete time (Caswell 2001). This multi-type branching process is not unique. The model used here to describe the population dynamics of *F. candida* is an age-structured matrix model ***A*** (Eqn. S.1) with six age classes and parameter values which don’t change over time. The age classes comprise different life cycle stages of *Folsomia candida*. The duration of intervals between observations is assumed to be one week and ***A*** projects numbers in each age class from one week to the next, with individuals advancing one age class per week (age classes one to five). To permit the presence of individuals of any age while keeping the number of age classes limited, individuals of age class six reach this class after being censused in age class five and remain in the sixth age class from then on as long as they survive. The *i*,*j-*th element of matrix ***A*** (Eqn. S.1), where *i* denotes rows and *j* columns, equals the expected number of individuals of age class *i* contributed to the population state at (*t* + 1) by one individual in age class *j* present at the census of week *t*. These expectations depend on age-specific survival probabilities and fecundities. Age-specific survival probabilities (in the absence of culling) are denoted by *s*_*j*_ and age-specific fecundities by *f*_j_. Here, we use indices *j* to distinguish “departure” states in which individuals can be when they contribute to a future population state and indices *i* to distinguish “arrival” states where the individuals which they contribute are in. To examine the effects of culling, a culling parameter **γ** was added, which represents the probability per week that an individual or a clutch of eggs escapes a single instantaneous culling event at the end of that week. Observation times can always be shifted such that culling occurs right before census. Each individual has an equal probability of being predated, this factor **γ** is the same across all age classes and affects all transitions equally (Eqn. S.1). We assume that ***A*** is primitive such that the Perron-Frobenius theorem holds (Horn & Johnson 2013).

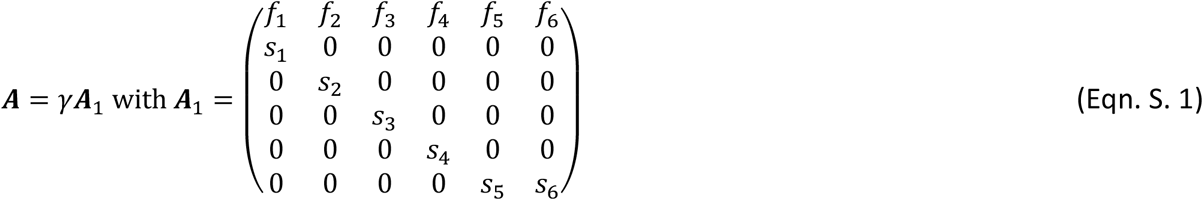

A large population where demographic stochasticity can be ignored and with its dynamics described by this model, will grow in size when the real and positive dominant eigenvalue **γ** of matrix ***A*** is larger than one, and will decrease in size when this eigenvalue is below one (Caswell 2001). Equivalently, one can calculate the expected lifetime reproductive success, which equals 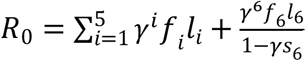, with 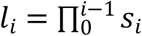 and *s*_0_=1. When *R*_0_ is larger than one, the eigenvalue **γ** of ***A*** also is (Cushing & Zhou 1994).

### Estimation of life history parameters *s* and *f*

We estimated the parameters of the life history model (Table 1) using approximations for birth-flow populations (Caswell, 2001), which account for the expected ages of individuals within an age class. Eggs of the clone “GM” hatch after a median development time of 9.5 days (Tully 2004). The fraction of eggs which hatch for this clone was estimated to be 0.989 (Tully 2023). This probability decreases for old individuals, but at older ages than the ones occurring in this experiment. Individual survival was estimated from hatched individuals, by observing them per hour (Tully 2023). An R script with our procedure to obtain parameter estimates of the matrix model is provided as supplementary material.

Age-specific survival probabilities per week were calculated based on an estimate of the baseline hazard per hour for six clones with similar life histories and a random “GM” clone effect. The baseline hazard was estimated using a kernel smoothed method (Mueller & Wang 1994) applied to all data on the pooled clones. We used the muhaz() function in R for this (https://cran.r-project.org/package=muhaz) with default settings, i.e., with a nearest neighbor bandwidth and boundary corrections. The random clone effect was estimated using a Cox model with frailty, i.e. a specific type of random effects survival model (Ripatti & Palmgren 2000) implemented in function coxme(), https://cran.r-project.org/web/packages/coxme/vignettes/coxme.pdf). The obtained values were interpolated to obtain cumulative hazards per hour. Survival across a certain interval after hatching is equal to 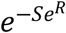 with *S* the hazard accumulated within that period and *R* the clone random effect reported by the mixed model output. Probabilities of survival before hatching for the clone “GM” were estimated based on the fraction of eggs which did not hatch. Given the low mortality of recently hatched individuals (Figure S. 1.), we assumed that death before hatching occurred within the first week of age and at a constant rate.

Following (Caswell 2001, eqn. 2.24 in there), we approximated the survival probabilities per age class as

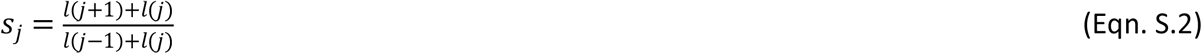

where *l*(*x*) is the probability of survival up to age *x*.

The minimum age at reproduction observed among the six clones was 299 hours (1.78 weeks) after hatching. A random effects ANOVA did not detect significant differences between clones in the first age at reproduction. When accounting for the egg development period, reproduction in the experiment started at age 3.14 weeks. For fecundities, we fitted the clutch sizes per hour per individual with zero-inflated mixed generalized linear Poisson models (mgcv library, Wood et al 2016) to the data on all six clones with similar life histories. These yield age-specific probabilities of producing a clutch per hour, and an age-specific clutch size conditional on producing a clutch in the concerned interval of one hour. In order to obtain fecundities per week for each age class, we predicted fecundities per hour using the generalized additive mixed model for clone “GM” and for all ages occurring within the first experiment. The prediction of the model at age 299 hours was a fecundity of value 0.031. We reset the values at earlier ages after hatching to zero. An approximation for birth-flow populations (eqn. 2.34 in Caswell, 2001) was applied to the resulting fecundities per hour to yield the *f*_j_.

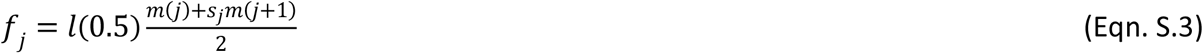

where *s*_*j*_is the survival probability in age class *j*, and where *m*(*j*) is summing all hourly predicted clutch size values from age *j* - 1 until (not including) age *j* in weeks. The resulting parameter values are listed in Table 1.

### Generating functions

We modeled the dynamics of small populations with finite size using discrete-time multi-type branching processes (Athreya & Ney 1972). The transpose of matrix ***A*** (Eqn. S.1) is a mean matrix (Athreya & Ney 1972) of different branching processes and contains parameters determining survival and offspring numbers shared with these stochastic processes. The dominant eigenvalue **γ** and *R*_0_ of ***A*** determine whether small populations have a chance at non-extinction, i.e. the probability that a branching process goes extinct after a finite number of generations is only smaller than one when **γ** and *R*_0_ are equal to or larger than one. In the so-called critical case (Athreya & Ney 1972) where **γ** and *R*_0_ are equal to one, the branching process always goes extinct ultimately when there is individual variation in the number of descendants. The supercritical case where **γ** and *R*_0_ are larger than one has a positive probability of non-extinction. However, these predictions do not state after how many time steps all extinctions are expected to have occurred, and how the probability of extinction changes with time. These depend on specificities of the branching process chosen to represent the generating process corresponding to matrix ***A*** (Caswell 2001, Jeppsson & Forslund, 2012)

Starting from a branching process, probabilities of extinction and establishment can be calculated using generating functions for the probability distributions of numbers of individuals, which, depending on what is calculated from them, are also called probability generating functions or reproduction generating functions (Jagers 1971), where reproduction can be loosely interpreted and mean to include survival of the individual itself (reproducing its presence).

We use the following notation (e.g., Sagitov & Shaimerdenova 2012) for vectors ***x*** (***y***) and vector arithmetic:

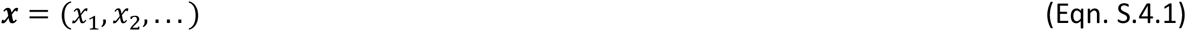

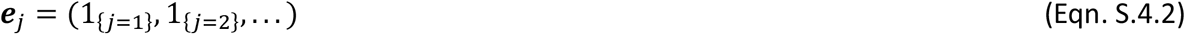

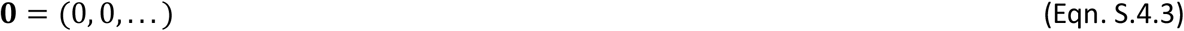

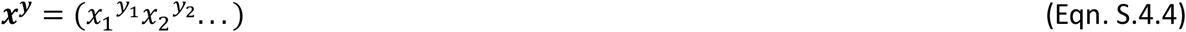

Where 1_{*j*=*k*}_ takes on value one for *j* = *k* and otherwise zero. Note that the result of expression (Eqn. S.4.4) is a scalar. The reproduction generating function of state or type *j* is defined as:

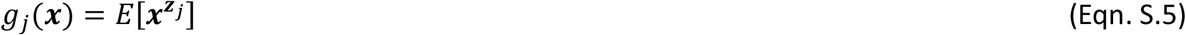

where ***x*** is the vector argument of the generating function which contains as many elements as there are states/types which have a non-zero probability to occur, with E[] denoting expectation and with vector ***z***_*j*_ = (*z*_1j_, *z*_2j_,…) containing the numbers of type *i* individuals produced by a type *j* individual across one time interval. Here, for the model of (Eqn. S.1), ***x*** and ***z***_*j*_ are *6*-tuples and ***x***^***z****j*^ =*x*_1_^*z*1*j*^*x*_2_^*z*2*j*^*x*_3_^*z*3*j*^*x*_4_^*z*4*j*^*x*_5_^*z*5*j*^*x*_6_^*z*6*j*^. We define the vector-valued generating function ***g*** as:

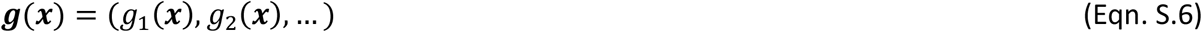

Using survival and fecundity parameters, a discrete time branching process can be defined by the probability generating function for the distribution of the number of descendants of each type of parent (Caswell 2001), including itself as a descendant, and which depends on the probability generating function of the number of offspring it will produce. Assuming that state transitions (here survival from one age class to another) and reproduction are independent, *g*_j_ is the product of generating functions for survival and for reproduction (Caswell 2001), which we will not treat explicitly here. The mean matrix of a branching process can be obtained by taking first derivatives of generating functions towards ***x*** (Athreya & Ney 1972). An easy way to obtain generating functions for a branching process from a matrix ***A*** seems to be multiplying each matrix cell by the corresponding argument of the generating function and adding the missing probability mass. However, one needs to specify the probability distributions of numbers of offspring. In order to arrive at a branching process and assuming for now that culling is not taking place, we assume that individuals in age-class *j* (*j* = 1 to 6) survive with probability *s*_j_ according a Bernoulli probability distribution. Individuals which can reproduce, produce a clutch each week, from which a random Poisson distributed number of juveniles *f*_*j*_ hatch the next week (Jeppsson & Forslund, 2012). The means of these Poisson distributions vary with age and are denoted by *f*_*j*_ (*j* = 1 to 6). Given the demographic parameters and their distributions, individuals reproduce independently which is a core assumption for branching processes. We will also assume that reproduction and transition to a different age class will occur independently within a time interval. When culling occurs once per week but not necessarily right before census, it will matter for the predictions of the branching process whether the first culling happens immediately after establishing or after one week. After that, if there is a positive probability to survive the first culling, the moment of observation of a process should not affect whether it is supercritical or not, as long as the process itself remains unchanged. Therefore, as long as culling events are spaced by one week, the specified process certainly goes extinct (eventually) if the dominant eigenvalue of matrix ***A*** (Eqn. S.1) is less than or equal to one. If this eigenvalue is larger than one, ultimate extinction occurs with a probability less than one and the population has a chance to persist indefinitely.

For age-structured multi-type branching processes, extinction probabilities vary as a function of the age classes of the individuals initially present in the population and the appropriate probability generating functions are therefore vector-valued (Bartlett 1951, Athreya & Ney 1972). We first derive branching process predictions for cases where culling occurs right before population census, and then propose how to derive predictions for different intervals between culling and observation and two possible culling per week.

We derive expressions for the six components of the vector-valued generating function for the branching process model we assumed and which corresponds to matrix model (Eqn. S.1). Even though we observed that eggs can be produced at any time, we assume for the derivation here that reproduction and survival before culling are statistically independent.

For a type *j* individual there are four different scenarios of events between censuses. (1) The individual as well as its clutch of eggs survive one time step. The probability that this happens equals *γ*^2^*s*_*j*_ and the result is the production of one individual of type *j* + 1 and a Poisson distributed number *z*_1*j*_ of individuals of type 1 in the clutch, with the expectation of this Poisson distribution equal *f*_j_. (2) The individual survives, but its clutch does not. This happens with probability *γs*_*j*_(1 − *γ*). (3) The individual dies, but its clutch survives. The probability of this scenario equal (1 − *γs*_*j*_)*γ*. (4) Neither the individual nor the clutch survives. The probability is (1 − *γs*_*j*_)(1 − *γ*). Thus, we obtain:

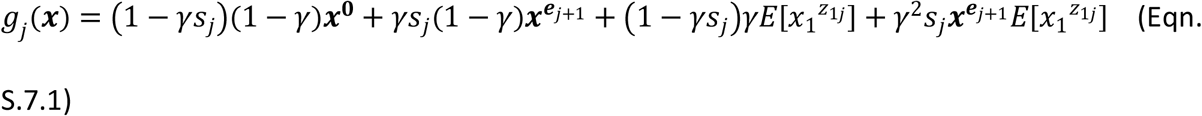

which becomes, using the generating function of the Poisson distribution 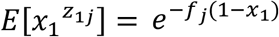 (e.g., Abramowitz & Stegun 1964),

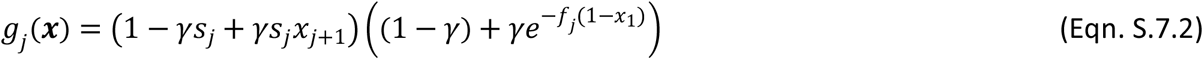

If there are no clutches produces by an age class *j* (*f*_j_ = 0), then this becomes

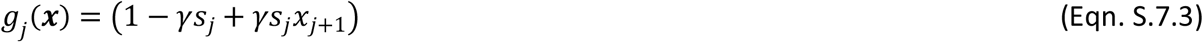

For age class six, *x*_*j*+1_ in Eqns. S.7.1-3 need to be replaced by *x*_*j*_.

### Interval subdivision

The branching process model above, was already tailored to the culling experiments carried out on *Folsomia*, but still had only a moderate amount of system-specific assumptions. It is possible to adapt the branching process model further to the experimental design and the biological system, in particular to heterogeneity within the interval between observations. When culling happens at a fixed time point during the interval between observations and not at the end of it, probabilities of outcomes can be affected by the interval length between observation and culling. Culling then divides an interval of length *t* between observations in two intervals of lengths *t*_1_ and *t*_2_ = *t* - *t*_1_. Probability generating functions can be defined for each separate interval, which together compose the probability generating function across the entire interval ***h***(***x***), such as is done for time-dependent branching rates (Dolgopyat et al. 2018). E.g.,

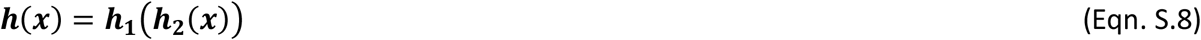

Where ***h***_**1**_ and ***h***_**2**_ are the probability generating functions for the numbers of individuals of different types produced across the first and second intervals, respectively. The vector ***x*** denotes the states which can be observed at census. These states don’t need to be the same at the end of each subinterval and therefore the vector-valued ***h***_**1**_ and ***h***_**2**_ can have vectors of different length as arguments. The number of components of ***h***_**1**_ is the same as of ***h*** and ***x***. The argument of ***h***_**1**_ has a length equal to the number of states observable at the end of the first subinterval and ***h***_**2**_ has the same number of components. When individuals reproduce either in the first or the second interval but not in both, then the number of states needs to increase to accommodate this memory effect. If entire clutches can be culled during an interval, we need to count numbers of clutches and not the amounts of eggs in them. With shorter intervals considered, individuals can hatch from some clutches only. We then need to differentiate those. The generating functions ***h***_**1**_ and ***h***_**2**_ are not specified separately, but were used to define ***h*** below with ***x*** and ***z*** which are ten-tuples.

It is now assumed that fecundity in age class one is zero. These individuals enter the age class as eggs, and are not expected to reproduce before they enter age class two as hatched individuals. Thus, unhatched individuals in different age classes are modelled as clutches (*z*_1_ to *z*_5_ are the numbers of clutches produced by age classes two to six), and the numbers of hatched individuals (five additional states with hatched individuals of age classes two to six *z*_6_ to *z*_10_). Clutches can be produced by individuals present at the start of intervals *t*_1_ and *t*_2_. An individual in state *j*, corresponding to age class *j* – 4, produces a clutch within *t*_1_ with probability *p*_j-4_, if not, it will do so in *t*_2_ when still alive. If the fecundity parameter for the parental age class is zero, *p*_j-4_ is always zero. It is assumed that clutches and individuals can be culled at the end of each subinterval, with probabilities *γ*_1_and *γ*_2_. Culling within an interval of an entire clutch already present at the start of an interval can occur. Survival of clutches is assumed to be equal to one in the absence of culling, of individuals in state *j* corresponding to age class *j* - 4 it is *s*_1,*j*−4_ and *s*_2,*j*−4_ in the two subintervals respectively.

A clutch produced by an individual of age class *k* in the previous time interval survives until after the first culling with probability *γ*_1_. It is assumed that all eggs from such a clutch hatch within the second interval and recruit to the second age class (which is the sixth state here). We assume that the number of hatched individuals surviving until after the second culling is Poisson distributed with mean *s*_1_*f*_*k*_*γ*_2_.

Therefore, for *j* in 1, …, 5,

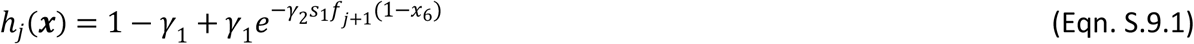

For a type *j* (j in 6, …, 10) hatched individual there are five different scenarios in the first subinterval. (1) The individual as well as its clutch, if produced, survive until after culling. The probability that this happens equals *p*_*j*−4_*γ*_1_^2^*s*_1,*j*−4_. (2) The individual reproduces and survives, but its clutch does not. This happens with probability *p*_*j*−4_*γ*_1_*s*_1,*j*−4_(1 − *γ*_1_). (3) The individual did not reproduce and survives. This happens with probability (1 − *p*_*j*−4_)*γ*_1_*s*_1,*j*−4_. (4) The individual dies, but its clutch survives. The probability of this scenario equal *p*_*j*−4_ (1 − *γ*_1_*s*_1,*j*−4_)*γ*_1_. (5) Neither the individual nor a clutch survives. The probability is (1 − *γ*_1_*s*_1,*j*−4_)(1 − *p*_*j*−4_ *γ* _1_). Note that for none of these scenarios, the individual changes state. Each of these scenarios leads to different scenarios for the second subinterval. For scenario (1) there are no individuals at the end of the second period. For (2), the individual dies with probability (1 − *γ*_2_*s*_2,*j*−4_)or survives and transitions to the next age class otherwise. Following (4), the clutch survives culling with probability *γ*_2_ and otherwise dies. For (3) and (5), with probability 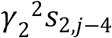 the individual as well as its clutch survive until after culling. The result is the production of one individual of type *j* + 1 and one clutch of type *j* - 5. With probability *γ*_2_*s*_2,*j*−4_(1 − *γ*_2_)the individual survives, but its clutch does not. With probability (1 − *γ*_2_*s*_2,*j*−4_)*γ*_2_ The individual dies, but its clutch survives. With probability is (1 − *γ*_2_*s*_2,*j*−4_)(1 − *γ*_2_)neither the individual nor the clutch survives.

Thus, we obtain for j six to nine:

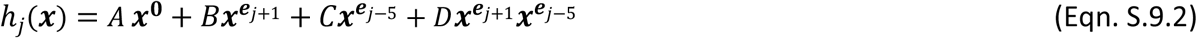

With

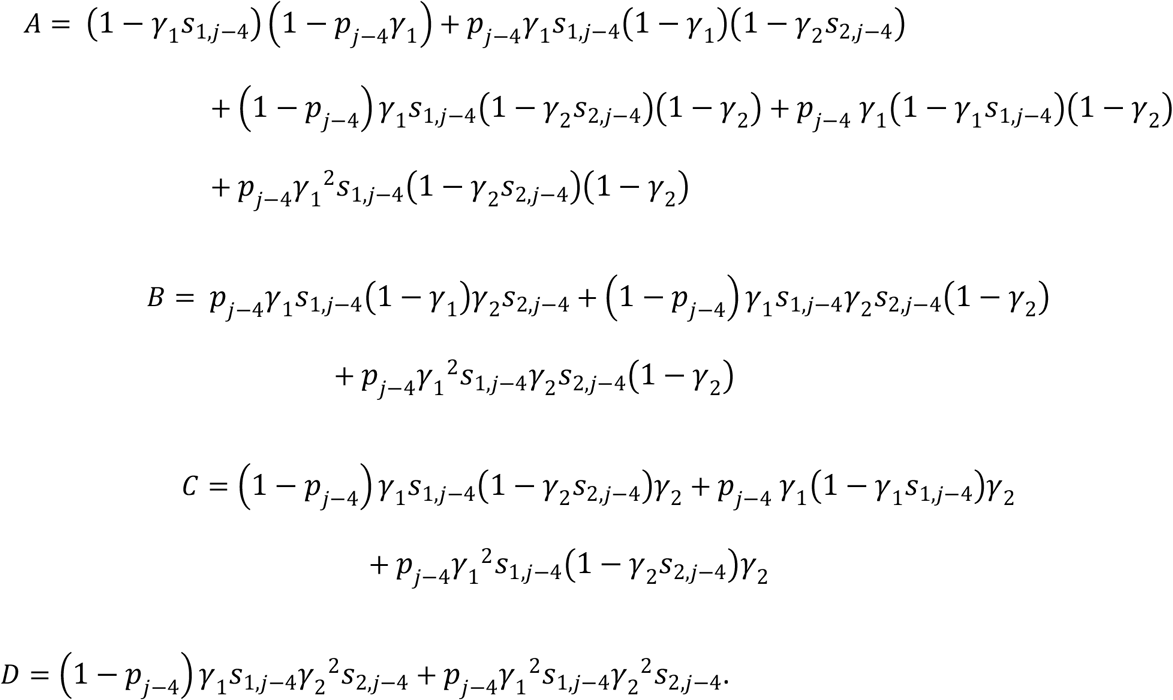

and 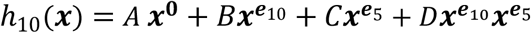. The mean matrix of this branching process can be derived using partial derivates of ***h***(***x***)(Athreya & Ney 1972). The matrix generally used in matrix population models is the transpose of this with elements 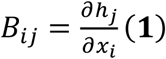. It can be found to be equal to

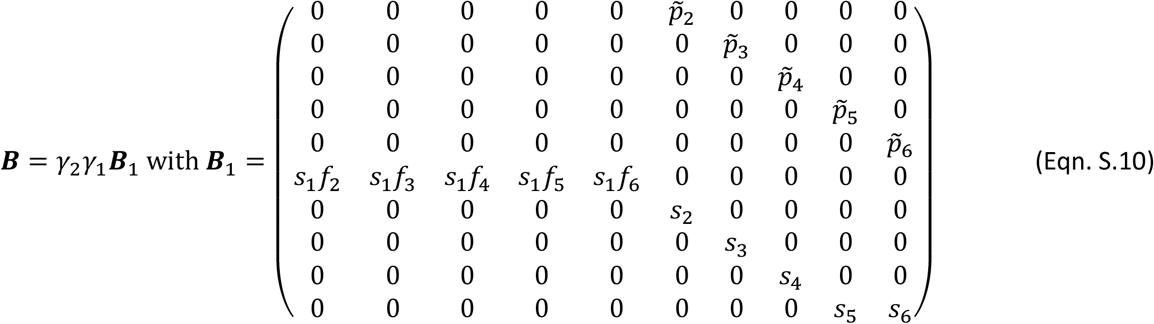

With 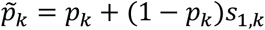. Denoting the sixth state as the birth state, lifetime reproductive success can be found to be equal to 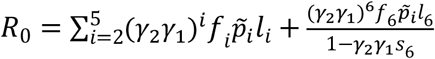, with 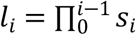 and *s*_0_ =1. Assuming that all *p* are equal to one and *γ*_1_*γ*_2_ =*γ*, one can find that 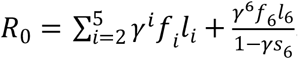. This is the same expression as above, when assuming that *f*_1_ is equal to zero.

### Survivorship functions

For practical purposes, such as comparing model predictions with empirical data, it is necessary to compute survivorship functions for populations initiated at time zero by a set of individuals with known ages. Survivorship functions give the expected fraction of these populations that survives a certain time span, given their initial composition. Here we show how to compute survivorship functions for populations initiated at time *t* = 0 from a single individual and monitored at weekly intervals (*t* = 0, 1, …., *n*, …). This is done using the generating functions (Eqns. S.6 and S.7) above.

For a population initiated from a single individual of type *j*, we denote the numbers of individuals in different states *i* in a population at time *t* (*t* a non-negative integer) by a vector ***Z***_*j*_(*t*), with elements *Z*_*ij*_(*t*) (*i* = 1, …, 6). The survivorship function for initial condition *j*, i.e., a population founded by one individual in state *j* at time zero is defined as:

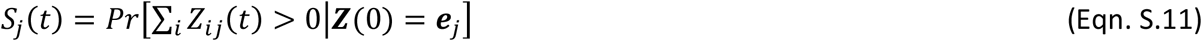

with ***Z***(0)the population state vector at time zero. Survivorship functions can be calculated by means of generating functions (Eqns. S.5 and S.6) with a recursive method (Athreya & Karlin 1971, Caswell 2001). The vector-valued generating function of the number of individuals ***Z***_*j*_(t) at time *t* is defined as:

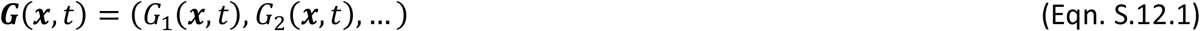

With

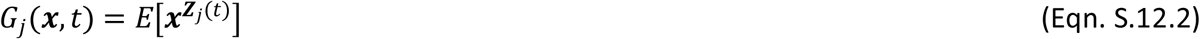

All individuals in ***Z***_*j*_(*t*)are contributed by individuals present at time *t* – 1. At each time step, the individuals present found from then on independently branching lines of descent. One can also see the individuals in ***Z***_*j*_(*t*)as the summed numbers of individuals in all descendant lines contributed by the individuals in ***Z***_*j*_(1). These individuals in ***Z***_*j*_(1)are in fact what we described as random variable ***z***_j_ above, the contribution of an age class *j* founder at the next census. The contributions of the descendants of each separate *i-*type individual in ***Z***_*j*_(1) = ***z***_*j*_ to the population state at time *t* is ***Z***_*i*_(*t* − 1), because the branching process is time-homogeneous. Using an expression for conditional multivariate generating functions (e.g., Bartlett 1951), one can thus obtain an equation to update ***G*** recursively, forward in time:

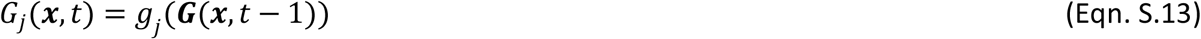

If we let *Q*_j_(*t*) denote the probability of extinction by time *t* of a population founded by a single type-*j* individual, it can be seen from (Eqn. S.9.2) that:

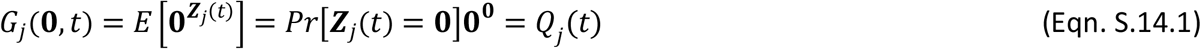

and that

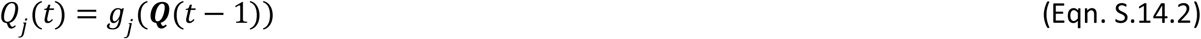

with ***Q*** the vector of extinction probabilities *Q*_j_ for populations founded by the respective types. This implies that we can use a forward recursive procedure to calculate the survivorship functions predicted by our model. The following system of equations (based on Eqns. S7.1-7.3), for *j* from one to six, holds for successive time steps and can be used to obtain extinction probabilities recursively:

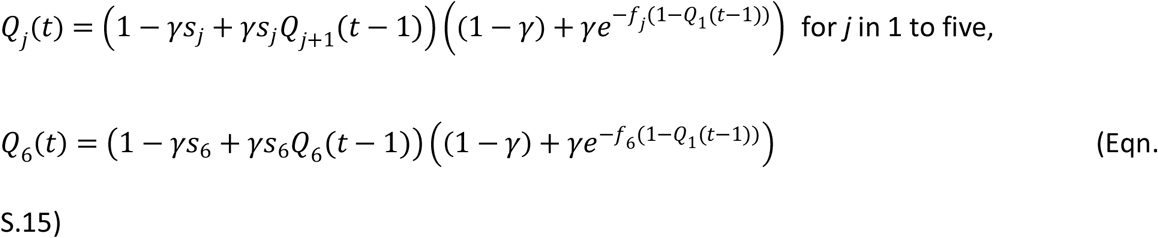

The initial probabilities *Q*_*j*_(0) are all equal to zero and populations founded with different individuals are extinct when none of the descendants of any founder is still alive, which is equal to the product of all individual extinction probabilities *Q*_*j*_(*t*)for the different founders. From the *Q*_j_(*t*) the survivorship functions for populations started with a single individual can be calculated as:

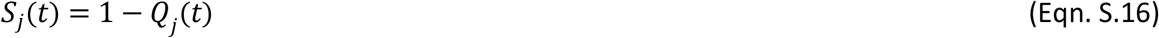

and the probability that a population at risk of extinction (non-extinct and not censored before time *t*) at time *t* - 1 goes extinct by time *t* equals:

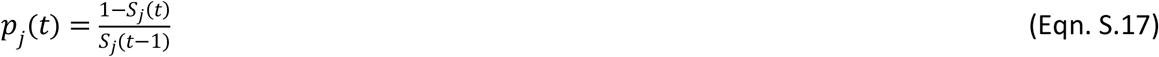

### Ultimate extinction probability

In multi-type branching processes, time-dependent extinction probabilities are expected to attain equilibrium values. After some time, all non-extinct populations are expected to contain that many individuals so that their conditional extinction probabilities after that time drop to zero. Let 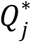 denote these equilibrium probabilities that a population goes extinct if we start with one individual of type *j*. For model (Eqn. 7), they can be calculated by solving the set of equations (*j* = 1, …, 6),

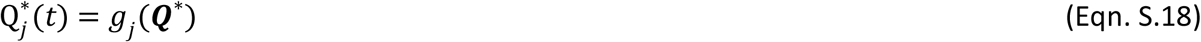

When the dominant eigenvalue of ***A*** (Eqn. S.1) is larger than one, this set of equations has a real non-negative solution with all 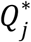 less than one, which corresponds to the vector of extinction probabilities for populations initiated with a single individual of a specific type. Numerical methods can be used to solve (Eqn. S.18). In fact, for the model of (Eqn. 7), one only needs to find 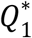 and the system of equations in (Eqn. S.15) will provide all other probabilities. For the model (Eqn. S.9), ***g*** was replaced by ***h*** and forward iteration was used to obtain the equilibrium ultimate extinction probabilities. A script calculating the survivorship functions and extinction probabilities was coded in R and is provided as Supplementary Material.

## Conditional mean extinction times

Experiments have a finite duration, and therefore only extinctions can be observed which occurred up to a certain time horizon *t*. Here we derive the expected extinction time conditional on this time horizon. Let *t* denote the extinction time, i.e., the time when an extinction is observed, then the conditional mean extinction time up to time *t* for a population founded by a single individual in age class *j*, *µ*_τ*j*_(*t*), equals:

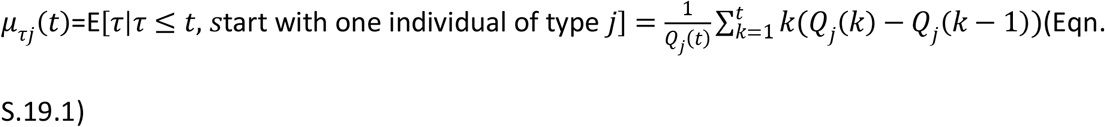

which is equal to:

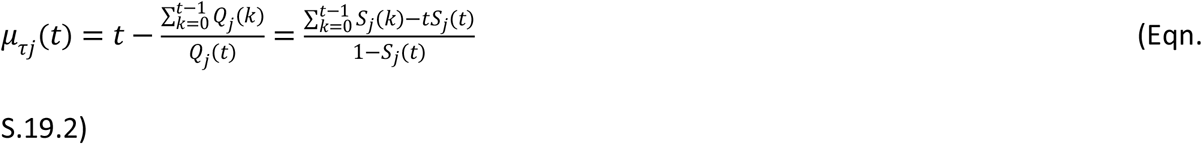

The supplementary R script provided calculates these conditional mean extinction times.

### Sensitivity analysis

Standard expressions (Caswell 2001) can be used to calculate the sensitivity of population growth rate to parameters in the matrix model ***A*** (Eqn. S.1). It can be shown that 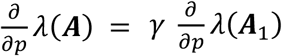 where *p* can be any of the parameter *s* or *f*. For the sensitivity to *γ*, 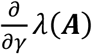, we observe that it is independent of the value of *γ* and a weighted sum of sensitivities of λ(***A***_1_)to the values of each of the matrix elements, where the weights are fecundity and survival parameters. The resulting values are given in Table 1, for λ(***A***_1_).

We investigated the sensitivity of extinction probabilities 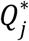 to the parameters in the matrix model ***A*** (Eqn. S.1). One parameter was varied at a time, keeping all other fixed. We estimated sensitivities of the eventual extinction probabilities 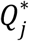 on survival parameters *s*_j_, on all age-dependent fecundities *f*_j_ and on *γ*. These are thirteen parameters in total. To determine the sensitivities to each parameter, we determined one or two slopes of linear least-squares regressions of extinction probabilities on sequences of parameter values. These sequences consisted (1) of the original value *p* and 1, 2, 3 and 4% increments of that value and (2) 1, 2, 3 and 4 % decrements. For fecundities we only considered increments and did not use sequence (2), for survival probabilities we used sequence (1).

## Figures

**Figure S.1.**
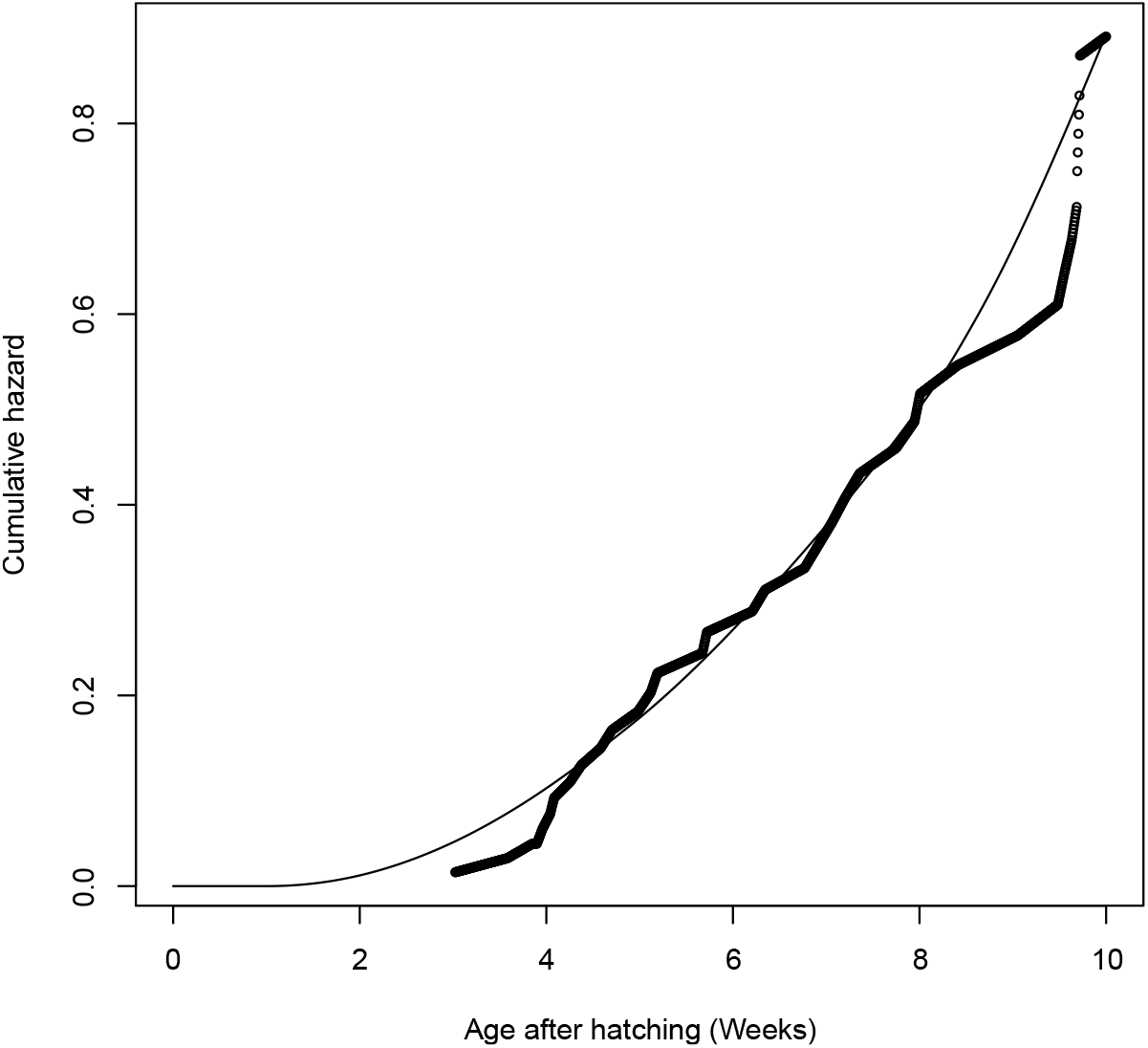
Cumulative baseline mortality rate after hatching. Cumulative mortality rates or hazards were estimated using a kernel smoothed method applied to all data of the five clones, interpolated to a value per hour. The result is plotted as a line, with age expressed in weeks. Nelson-Aalen estimates of the cumulative hazard are added as points for comparison.

**Figure S.2.**
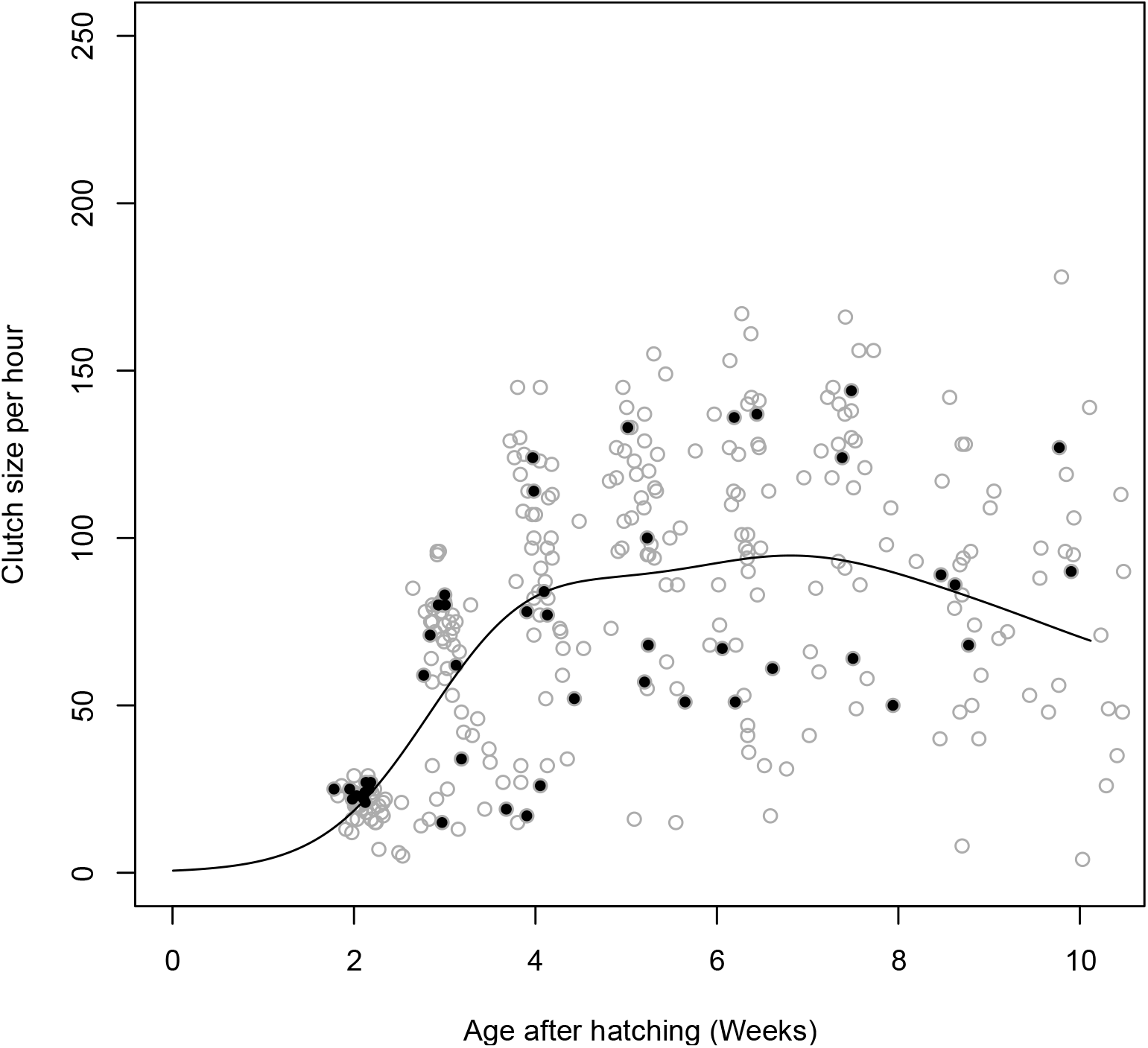
Observed and predicted rates of reproduction. Clutch sizes are plotted for all six clones with similar life histories observed by Tully (2004, 2023), data reproduced with permission. Clutch sizes for the clone “GM” are plotted in black. The prediction of age-dependent clutch sizes given that reproduction occurs and based on a zero-inflated Poisson model for the “GM” clone is superposed as a black line.

